# Comparative transcriptomic rhythms in the mouse and human prefrontal cortex

**DOI:** 10.1101/2024.09.26.615154

**Authors:** Jennifer N. Burns, Aaron K. Jenkins, Xiangning Xue, Kaitlyn A. Petersen, Kyle D. Ketchesin, Megan S. Perez, Chelsea A. Vadnie, Madeline R. Scott, Marianne L. Seney, George C. Tseng, Colleen A. McClung

## Abstract

Alterations in multiple subregions of the human prefrontal cortex (PFC) have been heavily implicated in psychiatric diseases. Moreover, emerging evidence suggests that circadian rhythms in gene expression are present across the brain, including in the PFC, and that these rhythms are altered in disease. However, investigation into the potential circadian mechanisms underlying these diseases in animal models must contend with the fact that the human PFC is highly evolved and specialized relative to that of rodents. Here, we use RNA sequencing to lay the groundwork for translational studies of molecular rhythms through a sex-specific, cross species comparison of transcriptomic rhythms between the mouse medial PFC (mPFC) and two subregions of the human PFC, the anterior cingulate cortex (ACC) and the dorsolateral PFC (DLPFC). We find that while circadian rhythm signaling is conserved across species and subregions, there is a phase shift in the expression of core clock genes between the mouse mPFC and human PFC subregions that differs by sex. Furthermore, we find that the identity of rhythmic transcripts is largely unique between the mouse mPFC and human PFC subregions, with the most overlap (20%, 236 transcripts) between the mouse mPFC and the human ACC in females. Nevertheless, we find that basic biological processes are enriched for rhythmic transcripts across species, with key differences between regions and sexes. Together, this work highlights both the evolutionary conservation of transcriptomic rhythms and the advancement of the human PFC, underscoring the importance of considering cross-species differences when using animal models.

## Introduction

The human prefrontal cortex (PFC) is a complex structure associated with many functions, including higher order cognitive function, goal directed behavior, and emotional regulation [1]. Two regions of particular interest are the dorsolateral prefrontal cortex (DLPFC), which is associated with cognition and plays a critical role in working memory, and the anterior cingulate cortex (ACC), which is implicated in both emotional and cognitive functions [1–3]. Notably, alterations in the DLPFC and the ACC have been linked to psychiatric diseases such as schizophrenia and major depressive disorder (MDD) [3–5].

To investigate the role of the PFC in the pathophysiology of psychiatric diseases, many studies utilize mouse models. However, there is widespread debate about the cross-species homology between the human and mouse PFC [6]. Indeed, while similar functions are attributed to both the mouse medial PFC (mPFC) and the human DLPFC, such as delay period activity during a working memory task [7], the cytoarchitecture of the mouse mPFC most closely resembles that of the human ACC. Specifically, both the mouse mPFC and the human ACC lack a granular layer 4 [6]. Likewise, studies have suggested a role of the rodent mPFC in conflict monitoring, reminiscent of findings in the human ACC [8,9]. Therefore, it remains unlikely that the mouse mPFC fully encapsulates the functions of a single human PFC subregion; instead, it likely represents a spectrum of features associated with different subregions of the highly diversified human PFC.

Circadian rhythms are ∼24-hour rhythms present in biological processes. These rhythms are produced through the molecular clock, a transcriptional-translational feedback loop (TTFL) consisting of CLOCK and BMAL1, which dimerize and promote the transcription of *Per* and *Cry*. PER and CRY then feedback and inhibit the activity of CLOCK and BMAL1, creating an ∼24 hour cycle of gene expression [10]. Beyond the TTFL, core clock genes bind to E-box elements, generating rhythms in the expression of up to 80% of protein coding genes [11,12]. Interestingly, polymorphisms in core clock genes as well as changes in rhythms have been linked to multiple psychiatric diseases and are thought to represent a key feature of disease pathophysiology [13–15]. Indeed, recent studies using human postmortem tissue have shown that there are broad changes in transcriptomic rhythms in the PFC of people with psychiatric diseases, including dampened rhythms in both the DLPFC and ACC of individuals with MDD and widespread circadian reprogramming in the DLPFC in schizophrenia [16,17]. As daily rhythms are present in functions associated with the PFC [18,19], alterations in transcriptomic rhythms may contribute to broad PFC dysfunction in psychiatric disease.

In this study, we use RNA sequencing to compare rhythms in the transcriptome of the mouse mPFC to two psychiatric disease relevant subregions of the human PFC, the DLPFC and the ACC. Our results indicate that rhythms in core molecular clock components and circadian rhythm signaling are broadly conserved. We additionally uncover species, sex, and subregion differences in the identity and associated biological functions of rhythmic transcripts.

## Methods

### Mouse tissue collection and RNA extraction

Adult male and female C57BL/6J (Jax ID: 000664) mice were group housed under a 12:12 light-dark cycle (lights on 0700, zeitgeber time (ZT)0, lights off 1900 (ZT12) with access to food and water *ad libitum*. All experiments were performed in compliance with University of Pittsburgh Institutional Animal Care and Use Committee guidelines.

Mice were sacrificed via cervical dislocation at 4-hour intervals across 24 hours (ZT2, 6, 10, 14, 18, 22); brains were removed and immediately placed on dry ice. Tissue punches of the mPFC (containing the anterior cingulate and prelimbic cortices) were taken from 150µm coronal sections cut on a cryostat (Leica Biosystems, Wetzlar, Germany). RNA was extracted using a RNeasy Plus Micro kit (Qiagen, Hilden, Germany).

### Mouse sequencing and data processing

Samples were assessed for concentration (Qubit, Thermo Fisher Scientific, Waltham, MA, USA) and RNA integrity (average: 9.1, standard deviation: 0.48) (Agilent RNA 6000 Kit, Agilent, Santa Clara, CA, USA). Library preparation was performed using a SMART Stranded Total RNA kit (Takara, Kusatsu, Japan) and samples were sequenced (2×101bp; 40 million reads/sample) using a Nova-Seq S4 (Illumina, San Diego, CA, USA). One sample (female ZT10) was excluded for failure to generate a library, leaving 4-5 mice/sex/timepoint for downstream analysis. Samples were evaluated for read quality using FASTQC and the per base sequence quality was high (average >34). Reads were aligned to the mouse reference genome (*Mus musculus* Ensembl GRCm38), converted to expression count data (HTSeq), and transformed to log_2_CPM (CPM=counts per million). Information on read counts can be found in Table S1. Counts were filtered with additional mouse mPFC sequencing samples and transcripts that did not meet a criterion of log_2_CPM>1 in >50% of the samples in at least one group, as well as genes on the Y-chromosome, were removed. Raw counts of the 13102 transcripts that met the filtering criteria were normalized by the median of ratios method in DESeq2 and log_2_ transformed [20]. UMAP plots were created to examine the clustering of mouse mPFC samples and additional mouse samples sequenced alongside the mPFC samples (Figure S1), as well as the clustering of mPFC samples by sex (Figure S2). To identify transcriptomic rhythms, we utilized a parametric cosinor model [21], whereby gene expression over a 24-hour period was fitted to a sinusoidal curve. R^2^ values were calculated as a measure of goodness of fit. P-values and FDR-corrected q-values were calculated from the F-test and used to determine rhythmicity.

### Human Samples

Sequencing data for the human samples were obtained from the CommonMind Consortium. All subjects had a known time of death, were <65 years old, and had a postmortem interval (PMI) of <35 hours. This cohort has previously been described in detail [22]. Briefly, subjects were matched between sexes within subregion for age, PMI, RNA integrity, and cause of death. There were no significant differences in these variables between sexes. In total, 84 subjects (42 per sex) were used for analysis of the DLPFC and 76 subjects (38 per sex) were used for analysis of the ACC. Transcripts were filtered based on the criteria outlined above and Y-chromosome and unidentified transcripts were removed, leaving 15239 transcripts for downstream analysis. Rhythmicity analysis was performed as described in [22].

### Comparison of rhythms and downstream analysis

Due to the exploratory nature of this study and limited sample sizes, transcripts with a p<0.05 were considered rhythmic and were used for downstream analysis. This is consistent with previous studies examining transcriptomic rhythms in the human and mouse brain [16,17,23,24]. We also utilized a threshold-free approach (rank-rank hypergeometric overlap (RRHO)) to assess differences in rhythmicity across groups [25]. Here, transcripts are ordered along the axes by -log_10_(p-value) and a heatmap is generated to visualize the overlap of rhythmic transcripts. To compare the phase shift in the timing of conserved canonical circadian genes (*PER1-3, DBP, CIART, NR1D1*) across species by sex, differences in the peak times of these transcripts between the human PFC subregions and the mouse PFC were pooled within sex and compared across sexes using a random effects model in the metafor package (R software). Ingenuity Pathway Analysis (IPA) (Qiagen) was used to assess functional pathways associated with rhythmic transcripts (p<0.05); a user-supplied background list of all transcripts meeting our filtering criteria was used for each analysis. Pathways that contained fewer than 15 genes were excluded. To compare between groups, the top 10 pathways enriched for rhythmic transcripts in each group were used to assess for significant enrichment in the opposite group. Pathways were considered enriched with a p<0.05 (-log_10_p-value>1.3).

### Data Availability Statement

Rhythmic transcripts in the mouse mPFC and human PFC subregions are listed in Tables S2-8. Sequencing data from the mouse mPFC will be shared to the Gene Expression Omnibus and sequencing data from the human samples can be obtained from the CommonMind Consortium.

## Results

### Transcripts in the mouse mPFC show diurnal rhythms in expression

In the mouse mPFC, we find that ∼12% of transcripts are rhythmic (1521 transcripts; p<0.05 cutoff) (Figure 1A) (Table S2). The top rhythmic transcripts, determined by p-value, include the core clock component *Arntl*, which encodes BMAL1, as well as transcripts such as *Ciart*, *Dbp*, and *Tef*, which have previously been shown to have conserved rhythmicity across tissues [12,26] (Figure 1B). Scatterplots showing the expression of the top two rhythmic transcripts over a 24-hour period, along with the fitted sinusoidal curve, are displayed in Figure 1C.

**Figure 1.**
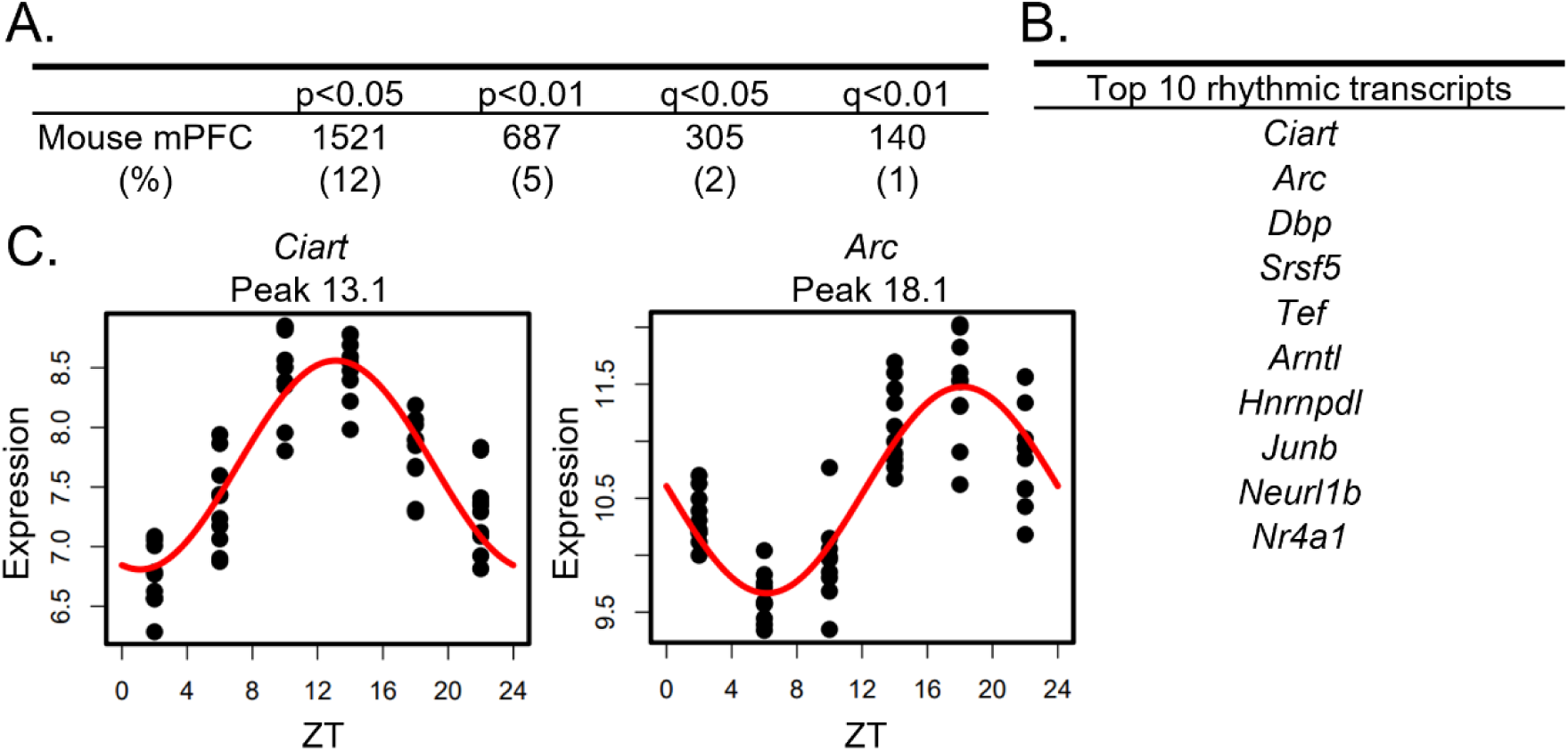
Rhythms in the mouse mPFC transcriptome. (A) The number (and percentage) of rhythmic transcripts detected in the mouse mPFC at different significance cutoffs. At a cutoff of p<0.05, 12% of transcripts in the mPFC are rhythmic. (B) The top 10 rhythmic transcripts in the mouse mPFC, as determined by p-value. Top rhythmic transcripts include known circadian genes including the core molecular clock component *Arntl*. (C) Scatterplots depicting rhythmic expression across 24 hours of the top two rhythmic transcripts. Time of death is depicted on the X-axis while expression is depicted on the Y-axis. Each point represents one subject. n=9-10/timepoint. mPFC=medial prefrontal cortex, ZT=zeitgeber time

### Rhythms in transcripts associated with the molecular clock are conserved across species

To compare transcriptomic rhythms between the mouse mPFC and human PFC subregions, we utilized a cohort containing data from the DLPFC and the ACC that has been previously described [22]. As this study found extensive sex differences in rhythmicity within the DLPFC and ACC, we performed our cross-species analysis stratified by sex. There are eight transcripts with conserved rhythms in the mouse mPFC and human PFC subregions of both sexes: *CHRM4, CIART, DBP, KANSL3, NR1D1, PER1, PER2,* and *PER3.* The majority of these transcripts (6/8) are associated with the molecular clock either directly (*PER1-3*) or through auxiliary loops that help to regulate the TTFL (*CIART, DBP,* and *NR1D1*) [10,27] (Figure 2). This is consistent with previous studies showing that rhythms in transcripts associated with the molecular clock are conserved between mice and humans in the brain [16].

**Figure 2.**
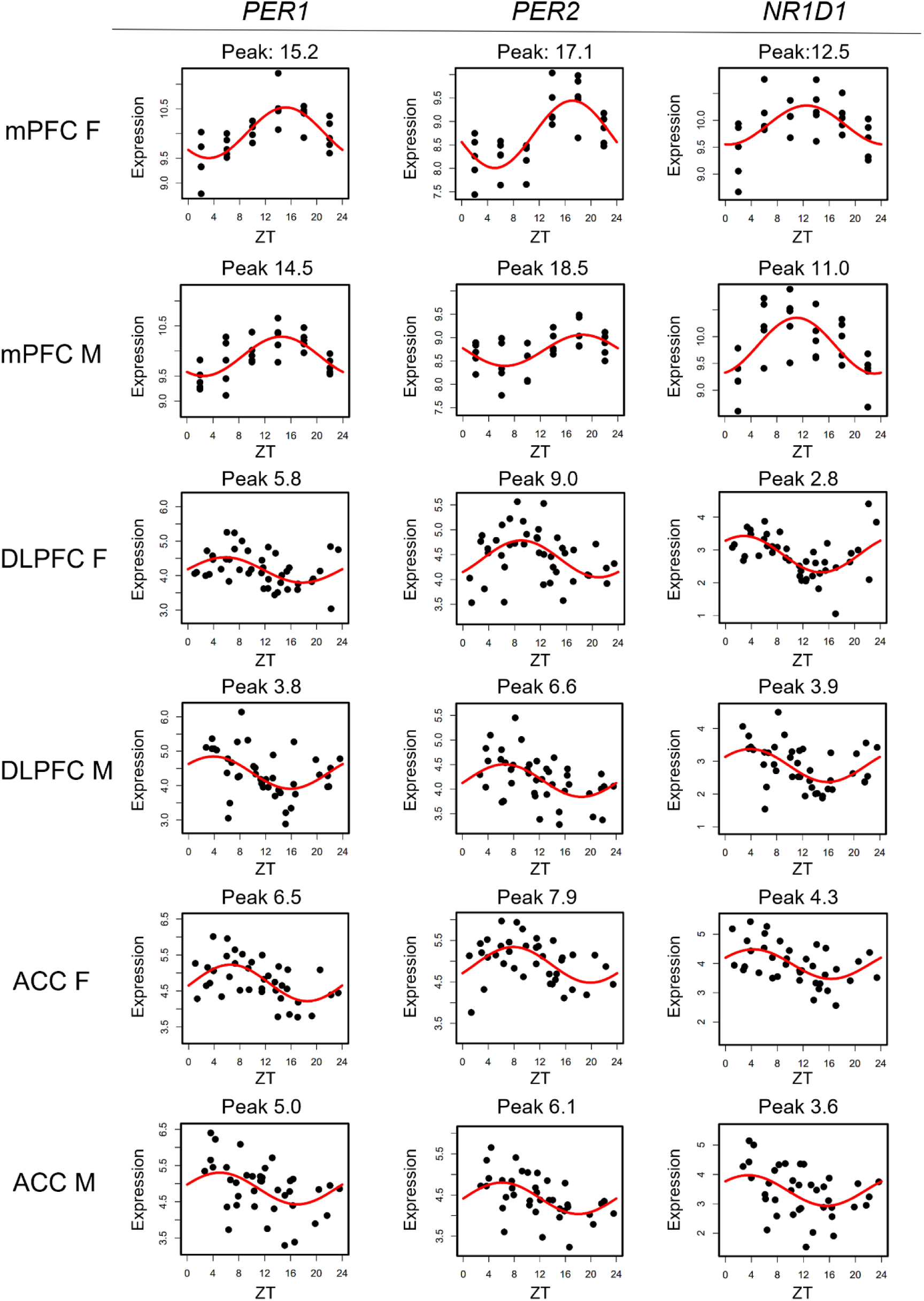
Rhythms in canonical circadian transcripts are conserved across species. Scatterplots depicting the expression of canonical circadian transcripts *PER1, PER2,* and *NR1D1* in the mouse mPFC and both human PFC subregions, separated by sex. Time of death is depicted on the X-axis while expression is depicted on the Y-axis. Each point represents one subject. mPFC=medial prefrontal cortex, DLPFC=dorsolateral prefrontal cortex, ACC=anterior cingulate cortex, ZT=zeitgeber time

### Peak times of canonical circadian transcripts differ across species by sex

We next determined how canonical circadian transcripts differ in their timing between the mouse mPFC and human PFC subregions (Table 1). We find a phase shift across species that differs by sex, with known circadian transcripts largely peaking ∼12 hours apart between the mouse mPFC and human PFC subregions in males. However, the difference in the peak time of canonical circadian transcripts across species is significantly smaller (p=0.002) in females, with most transcripts peaking only ∼9 hours apart. Moreover, previous studies have shown that in both the mouse suprachiasmatic nucleus (SCN) and the human PFC, *PER1* peaks first, followed by *PER3*, and then *PER2* [16,28], a pattern that we also observe across species in males. However, this temporal sequence is not observed in females in either the mouse mPFC or the human ACC. Therefore, while canonical circadian transcripts peak in opposing phases in the mouse mPFC and human PFC subregions, a finding that likely reflects differences in the active phase of each species, we also find sex differences in the timing of core clock components.

**Table 1.** Peak times of conserved transcripts differ across species by sex. The peak time (ZT) of all transcripts that are rhythmic in the mouse mPFC and both human PFC subregions in both sexes. The difference between the peak time in the human PFC subregion and the mouse mPFC of the respective sex is shown in parentheses. Broadly, transcripts peak closer in time across species in females than in males. mPFC=medial prefrontal cortex, DLPFC=dorsolateral prefrontal cortex, ACC=anterior cingulate cortex, ZT=zeitgeber time

|  | mPFC M | DLPFC M | ACC M | mPFC F | DLPFC F | ACC F |
| --- | --- | --- | --- | --- | --- | --- |
| *PER1* | 14.5 | 3.8 (-10.7) | 5 (-9.5) | 15.2 | 5.8 (-9.4) | 6.5 (-8.7) |
| *PER2* | 18.5 | 6.6 (-11.9) | 6.1 (-12.4) | 17.1 | 9 (-8.1) | 7.9 (-9.2) |
| *PER3* | 17.9 | 5.6 (-12.3) | 5.4 (-12.5) | 17.8 | 6.2 (-11.6) | 5.5 (-12.3) |
| *DBP* | 15 | 3.5 (-11.5) | 2.9 (-12.1) | 13.5 | 4.1 (-9.4) | 4.9 (-8.6) |
| *CIART* | 13.6 | 3 (-10.6) | 2.2 (-11.4) | 12.5 | 2.9 (-9.6) | 3.8 (-8.7) |
| *NR1D1* | 11 | 3.9 (-7.1) | 3.6 (-7.4) | 12.5 | 2.8 (-9.7) | 4.3 (-8.2) |

### Largest overlap in the rhythmic transcriptome between the mouse mPFC and ACC in females

To examine transcriptome-wide similarities, or differences, in rhythms between the mouse mPFC and human PFC subregions, we next assessed the degree of overlap in rhythmic transcripts across species. When comparing rhythmic transcripts (p<0.05) from each human PFC subregion to the mouse mPFC, we find that ∼5-10% of rhythmic transcripts are shared across species (Figure 3A), except for between the mouse mPFC and the ACC in females. Here, 20% (236 transcripts) of rhythmic transcripts in the mouse mPFC are also rhythmic in the human ACC. Notably, this overlap is not reciprocal, as only 8% of rhythmic transcripts in the human ACC are also rhythmic in the mouse mPFC in females. Using a more stringent significance threshold of p<0.01, we confirm that the greatest number of rhythmic transcripts (21 transcripts) overlaps between the mouse mPFC and human ACC in females (Fig. S3).

**Figure 3.**
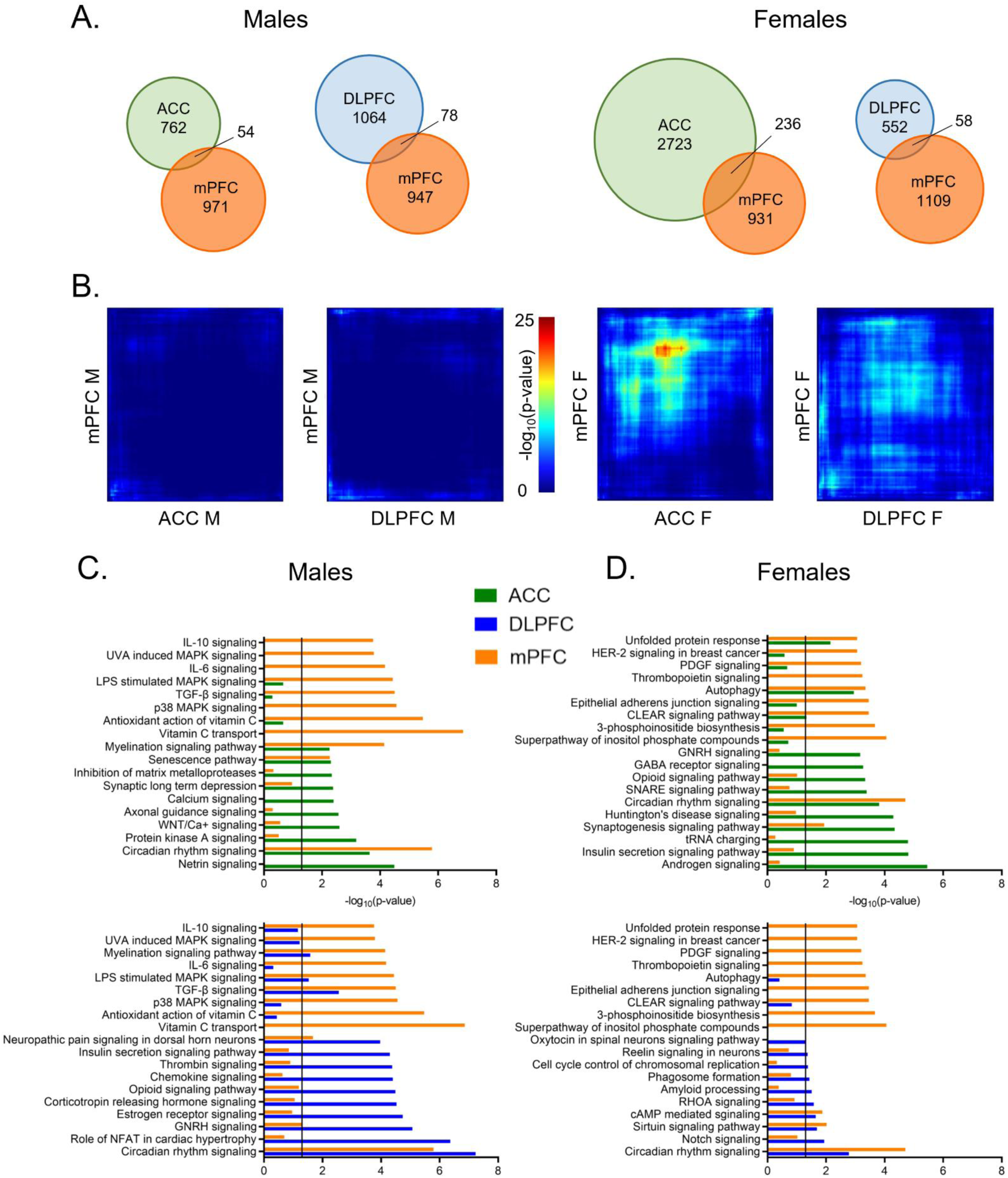
Greatest overlap in rhythmic transcripts between the mPFC and ACC in females. (A) Venn diagrams depicting the overlap in rhythmic transcripts (p<0.05) between the mouse mPFC and human PFC subregions, separated by sex. Twenty percent of rhythmic transcripts in the mouse mPFC are also rhythmic in the human ACC in females. All other comparisons across species share ∼5-10% of rhythmic transcripts. (B) Rank-rank hypergeometric overlap plots visualizing the overlap in rhythmic transcripts between the mouse mPFC and human PFC subregions. This threshold-free approach indicates that there is the most overlap in rhythmic transcripts between the mouse mPFC and the human ACC in females. (C) Ingenuity Pathway Analysis (IPA) was used to determine the top 10 pathways enriched for rhythmic transcripts (p<0.05) and their overlap between the mouse mPFC and human PFC subregions in males. (D) The top 10 pathways enriched for rhythmic transcripts (p<0.05), determined by IPA, and their overlap between the mouse mPFC and human PFC subregions in females. While the overlap in biological processes associated with rhythmic transcripts differs by region and sex, circadian rhythm signaling is among the top 10 pathways in the mouse mPFC and the human PFC subregions of both sexes. mPFC=medial prefrontal cortex, DLPFC=dorsolateral prefrontal cortex, ACC=anterior cingulate cortex

We next used rank-rank hypergeometric overlap (RRHO) plots as a threshold-free approach. Once more, we find that among all groups, there is the most overlap in rhythmic transcripts between the human ACC and the mouse mPFC from female subjects (Figure 3B). Rhythmic transcripts in the DLPFC of female subjects and the mPFC of female mice also show slight overlap, although less than that of the ACC. Notably, the RRHO analysis only uses transcripts that are expressed in both the mouse mPFC and human PFC subregions, suggesting that there are broad cross-species differences in rhythmicity even among the same subset of transcripts.

### Conservation of circadian rhythm signaling

Although the identity of rhythmic transcripts is largely distinct between the mouse mPFC and human PFC subregions, we next determined if the functions of rhythmic transcripts are conserved. Using Ingenuity Pathway Analysis (IPA), we find that circadian rhythm signaling is found among the top ten enriched pathways in all sexes, species, and subregions (Figure 3C&D).

In males, two additional pathways (senescence signaling and myelination signaling) are enriched for rhythmic transcripts in both the human ACC and the mouse mPFC (Figure 3C-top). Myelination signaling is also enriched for rhythmic transcripts in the DLPFC in males, suggesting a sex-specific conservation of rhythms in this process (Figure 3C-bottom). Additional pathways enriched in both the human DLPFC and the mouse mPFC of males include those associated with pain (neuropathic pain signaling in dorsal horn neurons), cytokine signaling (TGF-β signaling), and mitogen-activated protein (MAP) kinase signaling (LPS stimulated MAPK signaling). Many transcripts belonging to the MAP kinase family are also found within the enriched gonadotropin-releasing hormone signaling pathway. This suggests a conserved role of rhythms in intercellular signal transduction between the mouse mPFC and the human DLPFC in males.

We next performed the same analysis in females (Figure 3D). In addition to circadian rhythm signaling, four additional pathways are significantly enriched for rhythmic transcripts in both the human ACC and the mouse mPFC (Figure 3D-top). These pathways are associated with synaptogenesis, protein folding (unfolded protein response), autophagy, and lysosomal degradation (CLEAR signaling). When pathways enriched for rhythmic transcripts in the DLPFC and the mouse mPFC of female subjects are compared, we find less overlap in enriched pathways, consistent with our finding of less overlap in the identity of rhythmic transcripts (Figure 3D-bottom). In addition to circadian rhythm signaling, there are two pathways with significant overlap between the DLPFC and mPFC in females: sirtuin signaling and cAMP-mediated signaling. Many of the rhythmic transcripts that belong to the cAMP-mediated signaling pathway are G-protein coupled receptors (GPCRs), although the type of receptor differs between the human DLPFC and the mouse mPFC. Together, this indicates that although rhythmic transcripts differ in their identity, key biological processes, including neuronal signaling and protein processing, are enriched for rhythmic transcripts across species in females. However, in both sexes, over half of the enriched pathways are distinct between the mouse mPFC and human PFC subregions.

We next performed a cross species analysis of rhythms in the mouse mPFC and an evolutionarily intermediate species, using previously published data from male baboons [12]. While there are methodological differences between studies, we find that when compared to mice of the same sex, ∼20% of rhythmic transcripts in the mouse mPFC are also rhythmic in the baboon PFC (Fig. S4A). This is more than twice the proportion of rhythmic transcripts that are shared between the mouse mPFC and human PFC subregions in males. Moreover, there is greater overlap in the enriched biological processes (9 pathways) between the baboon PFC and mouse mPFC than between any of the human PFC subregions and the mouse mPFC (Fig. S4B). This suggests that there is greater conservation of transcriptomic rhythms between the mouse and baboon PFC than between the mouse and human PFC.

### Temporal patterns of rhythmic gene expression differ by species and sex

To examine overall patterns of rhythmic transcript expression across species, the percentage of rhythmic transcripts peaking at each timepoint (ZT) was plotted across 24 hours in two-hour bins. In males, nearly 60% of all rhythmic transcripts in the mouse mPFC peak during the active (dark) phase, with peak times fairly evenly distributed across the phase (Figure 4A). Similarly, in both human PFC subregions, the majority of rhythmic transcripts peak in the active (light) phase, with ∼68%, or over 80%, of rhythmic transcripts peaking during the active (light) phase in the human ACC and DLPFC, respectively.

**Figure 4.**
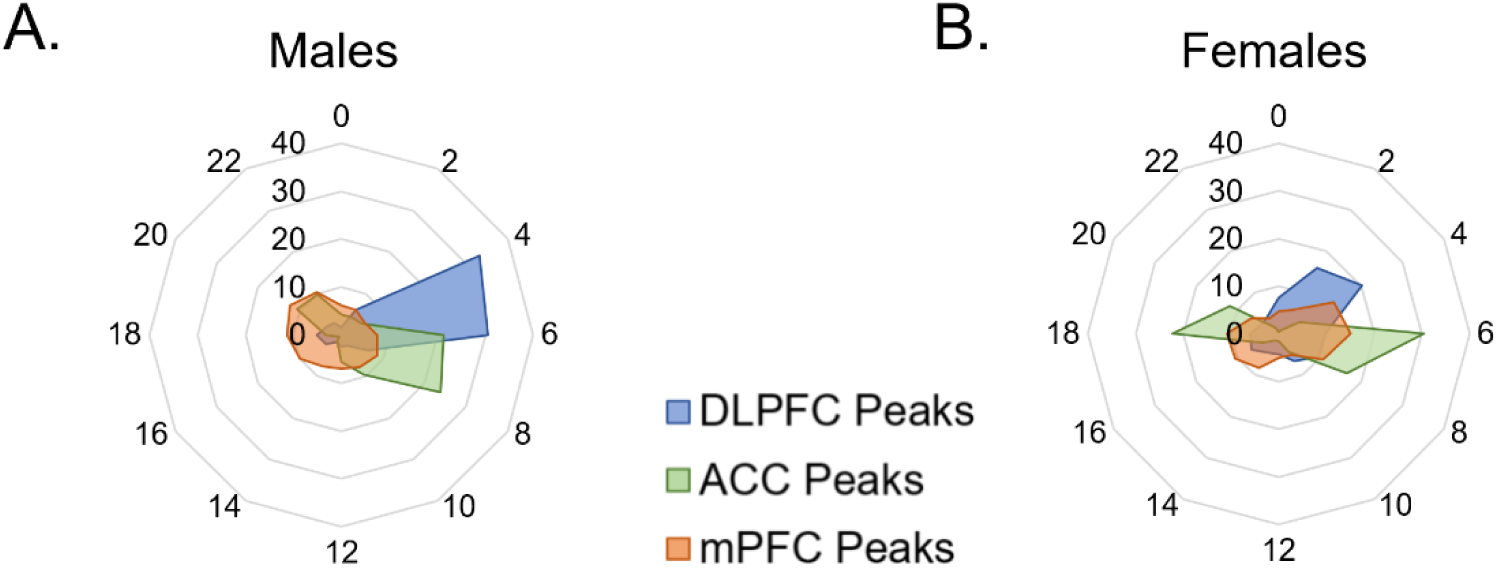
Temporal patterns of rhythmic expression vary by region, sex, and species. (A&B) The peak time of rhythmic transcripts (p<0.05), plotted as the percentage of total rhythmic transcripts peaking in each 2-hour bin across 24 hours (ZT) in (A) males and (B) females. In males, rhythmic transcripts largely peak in the opposite phase between the mouse mPFC and human PFC subregions. In females, over half of rhythmic transcripts peak in the active phase (light) in humans, whereas about half of transcripts peak in each phase in the mouse mPFC. mPFC=medial prefrontal cortex, DLPFC=dorsolateral prefrontal cortex, ACC=anterior cingulate cortex, ZT=zeitgeber time

In females, rhythmic transcripts in the mouse mPFC largely fall into two groups, with approximately half peaking in each phase (Figure 4B). Similarly, large groups of rhythmic transcripts in the human ACC peak in each phase, with ∼57% peaking in the active (light) phase and ∼43% peaking in the inactive (dark) phase. In contrast, in the human DLPFC of female subjects, most (∼70%) of the rhythmic transcripts peak during the active (light) phase. These data demonstrate that although the expression of core clock genes can be predicted by the active phase of the species, the temporal patterns of total rhythmic transcripts are highly variable between species, PFC subregions, and even sexes.

## Discussion

Recent literature has highlighted the importance of transcriptomic rhythms in brain health and disease [16,17,22]. In this study, we compared rhythms between the mouse mPFC and two subregions of the human PFC, the ACC and the DLPFC, that are heavily implicated in psychiatric disorders [3–5]. Consistent with previous studies, we find that canonical circadian genes are rhythmic in both humans and mice [16,26]. These conserved rhythmic transcripts largely peak in opposing phases in mice and humans, likely reflecting differences in the active phase of each species. However, we find that the difference in timing of core circadian genes between species depends on sex, with females showing a smaller shift than males (∼9 hours vs. ∼12 hours). This may be driven by sex differences in the rhythms of core clock genes within species. Indeed, previous studies found that the peak time of core clock gene expression differs by sex in the DLPFC of aged subjects [29], whereas studies in the rodent PFC found that rhythms in core clock genes were more robust in males [30]. While the mechanisms underlying sex differences in transcriptomic rhythms are currently unknown, we hypothesize that circulating hormones may have an effect. Indeed, the SCN expresses both estrogen and androgen receptors [31,32] and systemic estradiol administration has been shown to phase advance core clock gene expression in the SCN [31–33]. Moreover, one study found that in the PFC, rhythms in *Arntl* differed between rats with a normal estrous cycle and non-cycling rats [30]. Although differences in estrous phase/menstrual cycle were not assessed in this study, the proportion of rhythmic transcripts is not consistently lower in female subjects. Therefore, potential variability across the estrous/menstrual cycle likely does not impair our ability to detect rhythmic transcripts.

While most rhythmic transcripts that are broadly conserved across species and sexes are closely associated with the molecular clock, rhythms in two transcripts, *KANSL3* and *CHRM4*, are also broadly conserved. KANSL3, which is involved in chromatin remodeling, plays a role in regulating the transcription of housekeeping genes and facilitates the transcription of mitochondrial DNA in cells with high metabolic rates, such as neurons [34,35]. The conserved rhythmicity of this transcript suggests that molecular clock control over the transcription of housekeeping genes and mitochondrial function are evolutionarily conserved. On the other hand, *CHRM4* encodes the muscarinic acetylcholine receptor M4, which has been proposed as a therapeutic target for schizophrenia [36,37], Therefore, conservation of rhythms in *CHRM4* across species may be important for successful translation of these drugs into humans.

Nevertheless, we find that most of the rhythmic transcripts in the human PFC subregions are not rhythmic in the mouse mPFC of the same sex. Indeed, while 20% of rhythmic transcripts in the mouse mPFC are rhythmic in the ACC in females, only 8% of rhythmic transcripts in the human ACC are rhythmic in the mouse mPFC. This suggests that rhythms in gene expression changed as the human PFC evolved and became more specialized. Our findings are consistent with theories of PFC evolution, whereby agranular and dysgranular regions, such as the ACC, evolved earlier, while granular regions of the PFC, such as the DLPFC, evolved later and are considered to be unique to primates [38]. The idea that molecular rhythms diverged across evolution is further supported by our finding that there is greater overlap in rhythmic transcripts between the baboon PFC and the mouse mPFC, as well as findings from a recent paper showing that up to 38% of rhythmic transcripts are shared between mice and humans in the more evolutionarily conserved striatum [23]. Notably, however, we find that the elevated overlap between the mouse mPFC and the human ACC is specific to females, a finding perhaps driven by previously described sex differences in rhythmicity in the human ACC [22].

When the biological processes associated with rhythmic transcripts are assessed, we find that pathways associated with intercellular communication and the integration of extracellular signals are significantly enriched for rhythmic transcripts in both the mouse mPFC and human PFC subregions. The enrichment of rhythmic transcripts in these pathways suggests that control of the molecular clock over mechanisms associated with cellular signaling, including neurotransmission, are broadly conserved. Moreover, while the rhythmic transcripts belonging to each pathway are generally different between mouse and human, some are closely related. This is particularly true within the unfolded protein response pathway, whereby many rhythmic transcripts in both the human ACC and the mouse mPFC encode proteins in the DNAJ heat shock family.

Similar to previous studies, which found that the timing of total rhythmic transcripts varies widely even in anatomically adjacent tissues [12], we find broad differences in the temporal patterns of rhythmic gene expression across PFC subregions, sexes, and species. These patterns are much more variable than the timing of core clock genes, suggesting that they are generated downstream of the molecular clock. Indeed, studies have found that many targets of the molecular clock are transcription factors, resulting in rhythms in gene expression that are tissue specific [39]. It is likely that a similar mechanism underlies the differences found in this study. Understanding temporal patterns in gene expression, and what drives them, may have clinical implications for psychiatry. For example, a recent study found that the time in which antipsychotics were administered affected the development of metabolic side effects in both humans and mice [40]. However, the timing depended on the active phase of each species, highlighting the importance of considering cross-species differences in rhythms when translating preclinical findings into humans.

Differences in rhythms between the mouse mPFC and human PFC subregions may be partially attributable to cross-species differences in the cellular makeup of these regions. Indeed, the expression of markers used to define individual cell types differs across species in the cortex and studies have shown that, even within conserved cell types, gene expression differs between mice and humans [41,42]. Moreover, differences in the timing of transcripts across cell types may make transcripts appear non-rhythmic in homogenate tissue, as one study in the mouse SCN found that core clock genes peak earlier in neurons relative to non-neuronal cells [43]. It remains unknown whether similar differences in timing exist across cell types in the PFC and whether these differences are conserved across species.

Variation in the light-dark cycle may also contribute to the observed differences in rhythmicity between the mouse mPFC and human PFC subregions. While mice in this study were housed under a strict 12:12 light-dark cycle, the light-dark cycle of the human subjects was likely more variable. While the time of death of human subjects is normalized to sunrise, we cannot eliminate the effect of artificial light or differences in behavioral rhythms. Therefore, it is likely that increased variability in the human samples may result in an underestimation of true rhythmic transcripts. Undoubtedly, with higher sample sizes and more statistical power, we would identify additional rhythmic transcripts in humans.

As over 80% of proteins identified as druggable targets by the FDA show rhythms in gene expression [12], this study provides an important translational framework for understanding how these rhythms differ across species. These findings paint a complex picture, whereby molecular rhythms show distinct patterns based on sex, species, and PFC subregion, likely reflecting the unique functions of rhythmicity in the highly specialized human PFC. Given that patterns of rhythmic gene expression show extensive changes in the brains of individuals with psychiatric diseases, it is imperative to carefully consider differences in rhythms between species when mice are used for mechanistic studies into these disorders.

## Acknowledgements

We would like to thank the CommonMind Consortium for the data from the human postmortem tissue. We would also like to thank the University of Pittsburgh Health Sciences Sequencing Core and UPMC Genome Center for assistance with library preparation and RNA-sequencing.

## Author Contributions

Conceptualization: JNB, CAM; Methodology: GT, CAM; Data acquisition: JNB, AKJ, MLS, KAP, MSP, KDK, CAV; Data analysis: JBN, XX, KAP, MLS, GT, AKJ, KDK; Writing and editing: JNB, AKJ, XX, MRS, MLS, CAM,

## Competing interesting

The authors have nothing to disclose.

## Supplementary Figures

**Figure S1:**
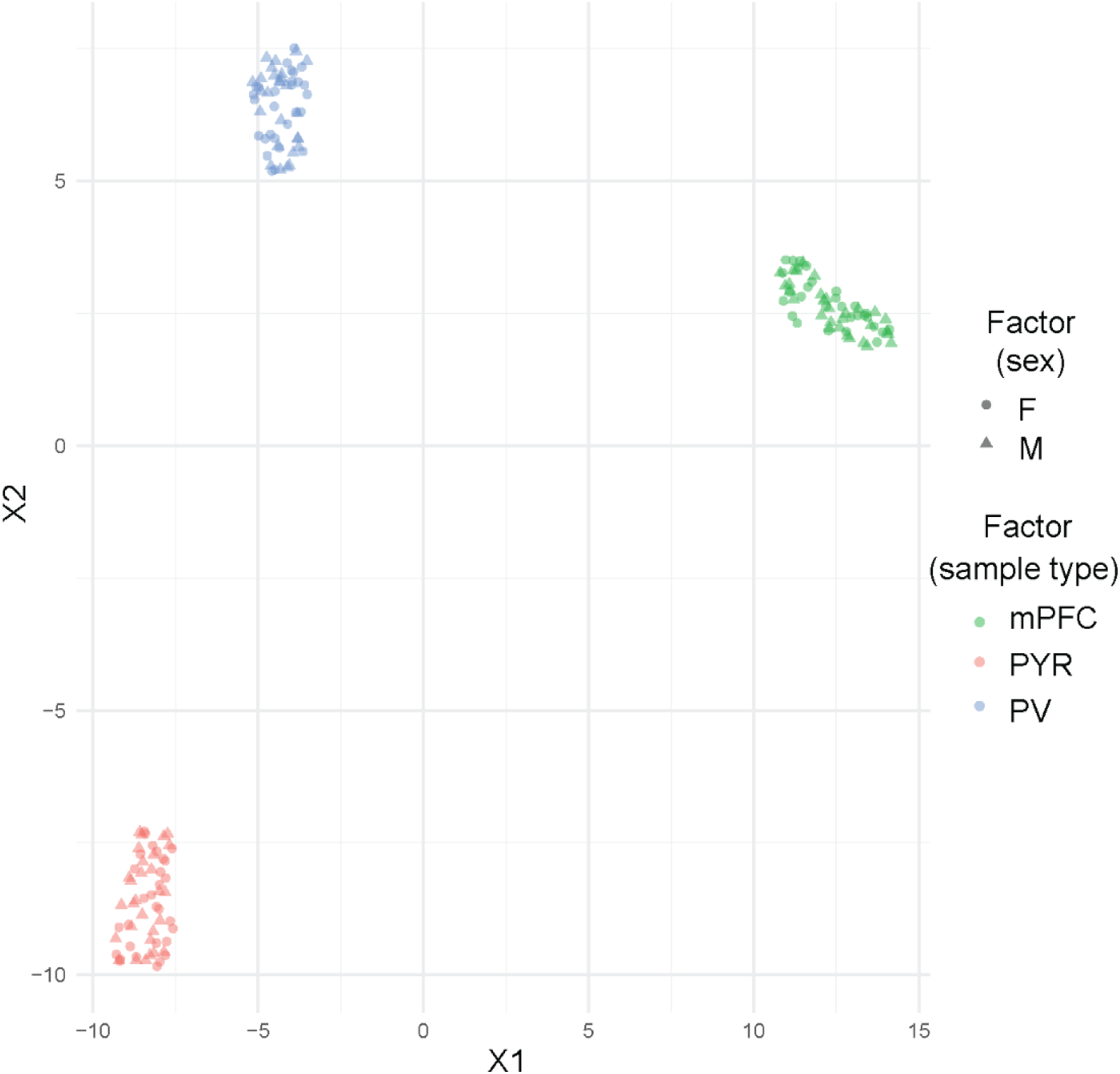
Samples form discrete clusters segregated by sample type. UMAP plot visualizing the clustering of three types of mouse samples sequenced together (homogenate mPFC and two cell-type specific samples-parvalbumin and pyramidal cells). mPFC=medial prefrontal cortex, PV=parvalbumin cells, PYR=pyramidal cells

**Figure S2:**
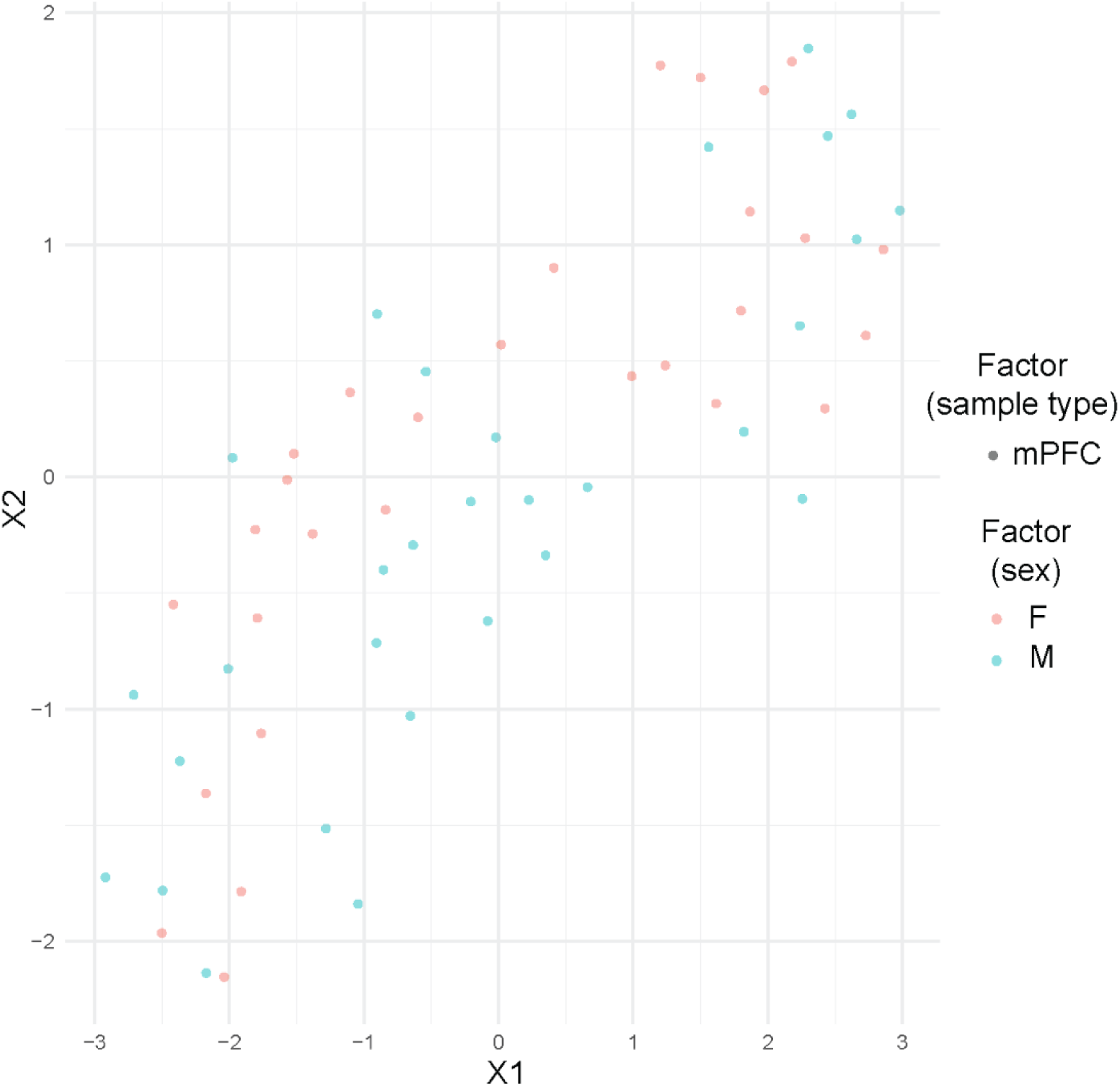
mPFC samples do not form transcriptionally distinct groups. UMAP plot visualizing the clustering of mPFC samples. mPFC=medial prefrontal cortex

**Figure S3:**
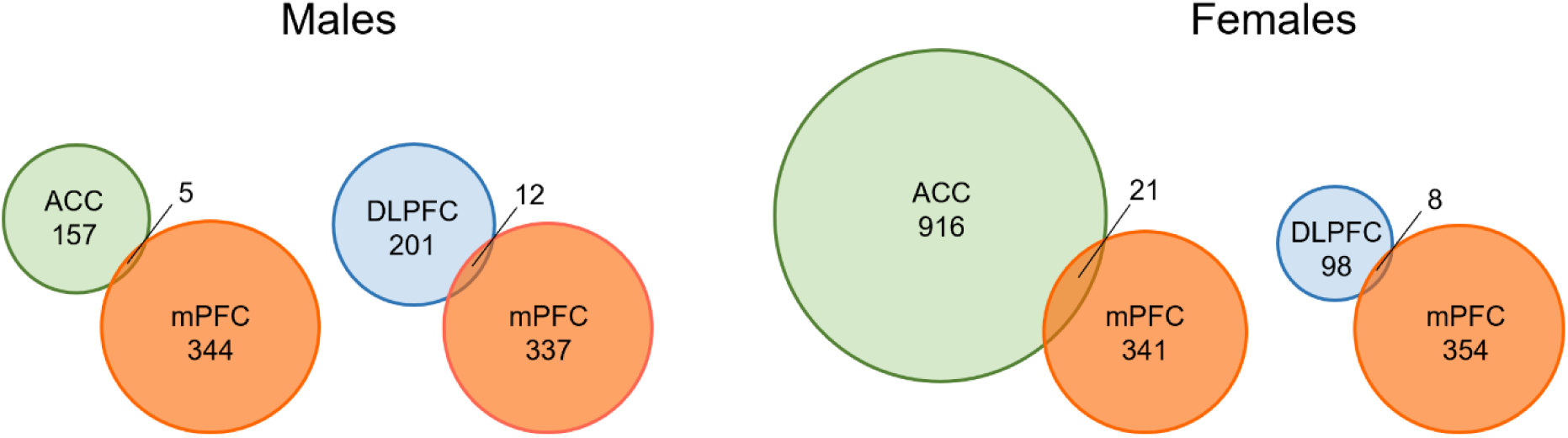
Greatest overlap in rhythmic transcripts between the mPFC and ACC in females at a more stringent cutoff. Venn diagrams depicting rhythmic transcript overlap (p<0.01) between the mouse mPFC and human PFC subregions. mPFC=medial prefrontal cortex, DLPFC=dorsolateral prefrontal cortex, ACC=anterior cingulate cortex

**Figure S4:**
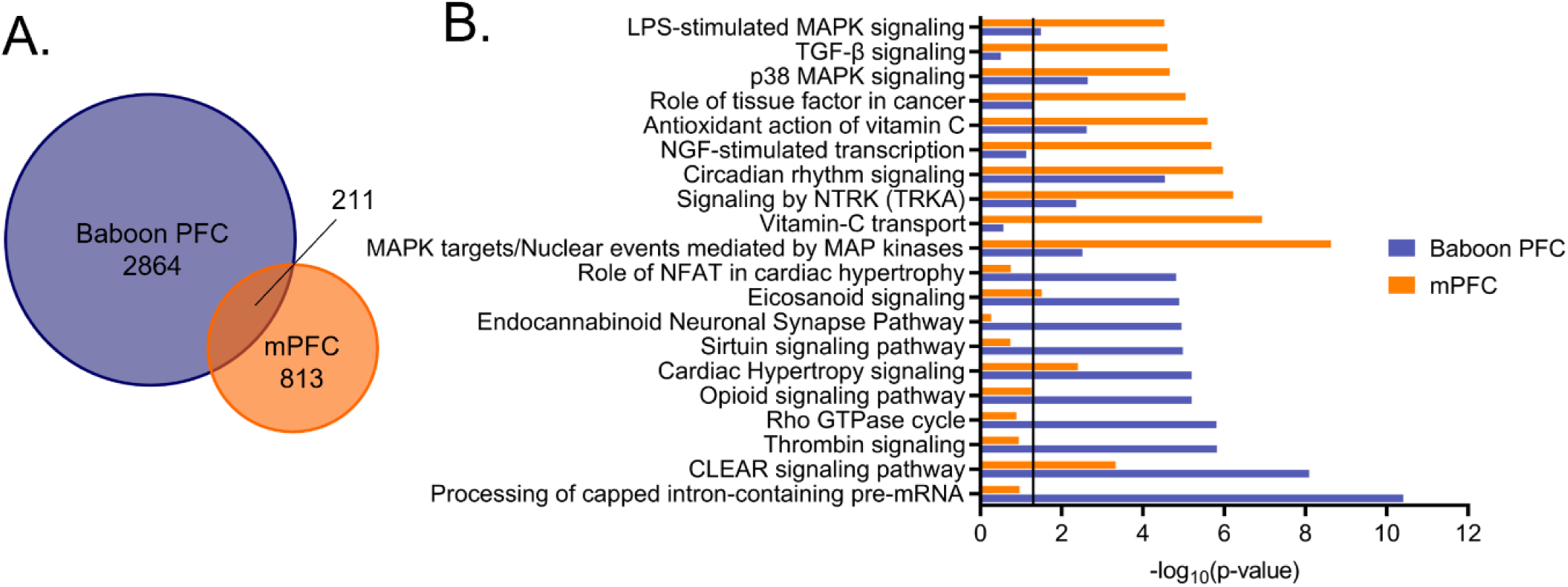
Greater conservation of rhythms between the mouse mPFC and the baboon PFC. (A) Venn diagrams depicting the overlap of rhythmic transcripts between the baboon PFC and mouse mPFC in males. Approximately 20% of transcripts that are rhythmic in the mouse mPFC are also rhythmic in the baboon PFC. (B) Overlap of the top pathways (identified by Ingenuity Pathway Analysis) enriched for rhythmic transcripts in the mouse mPFC and baboon PFC. Nearly half of the pathways (9) are significantly enriched for rhythmic transcripts in both the mouse mPFC and baboon PFC. Baboon data from [2]. mPFC=medial prefrontal cortex

## Supplementary Tables

Table S1: Total reads and read counts (total reads x alignment rate) for mouse samples

Table S2: Rhythmic transcripts (p<0.05) in mouse mPFC with sexes combined

Table S3: Rhythmic transcripts (p<0.05) in mouse mPFC in females

Table S4: Rhythmic transcripts (p<0.05) in mouse mPFC in males

Table S5: Rhythmic transcripts (p<0.05) in DLPFC in females

Table S6: Rhythmic transcripts (p<0.05) in DLPFC in males

Table S7: Rhythmic transcripts (p<0.05) in ACC in females

Table S8: Rhythmic transcripts (p<0.05) in ACC in males

